# Volume Electron Microscopy of Cortical Organoids: Methods for Region Identification, Connectome Reconstruction, and Organelle Segmentation

**DOI:** 10.64898/2025.12.31.697152

**Authors:** Sveva Dallere, Anna Mattioni, Marta Turégano-López, Lidia Blazquez-Llorca, Angel Merchàn-Perez, Roberta Schellino, Alessandro Vercelli, Javier DeFelipe, Marina Boido

**Affiliations:** Neuroscience Institute Cavalieri Ottolenghi, Department of Neuroscience “Rita Levi Montalcini”, University of Turin, Turin, Italy; University School for Advanced Studies IUSS Pavia, Pavia, Italy; Laboratorio Cajal de Circuitos Corticales, Centro de Tecnología Biomédica, Universidad Politécnica de Madrid, Pozuelo de Alarcón, Madrid, Spain; Department of Neurosciences and Biomedical Sciences, University Carlos III of Madrid, 28903 Madrid, Spain; Departamento de Tecnología Fotónica y Bioingeniería, ETSI Telecomunicación, Universidad Politécnica de Madrid, Madrid, Spain

**Keywords:** synapses, synapse segmentation, connectomics, CLEM, 3D tissue models, FIB/SEM, TEM

## Abstract

Volume electron microscopy (vEM) has become a powerful tool for 3D ultrastructural analysis of neural circuits, yet its application to human brain organoids remains limited, particularly for connectomic studies. Here, we established a comprehensive and scalable workflow for applying vEM to human cortical organoids, integrating correlative light and electron microscopy, large-area SEM mosaic imaging, focused ion beam–scanning electron microscopy (FIB-SEM), and transmission electron microscopy (TEM) validation.

By systematically comparing two embedding protocols in use, we demonstrated that the DeFelipe and Fairén (1993)/Cano-Astorga et al. (2024) method provides optimal compatibility with toluidine blue–stained semithin sectioning and enables reliable synapse segmentation and neurite tracing. In contrast, the Deerinck et al. (2010) protocol offers enhanced membrane contrast but limits postsynaptic density visualization. Using FIB-SEM imaging of peripheral, neuropil-like regions of cortical organoids, we achieved accurate 3D reconstruction of synapses, neurites and intracellular organelles, enabling quantitative assessment of synaptic apposition surfaces, neurite trajectories, and organelle distribution across defined cellular compartments.

Together, our results demonstrate for the first time the feasibility of micro-connectomic reconstruction in human cortical organoids at nanometer resolution. This methodological framework expands the applicability of vEM to organoid systems and provides a robust foundation for future studies of human brain development, disease modeling, and therapeutic evaluation at the synaptic and subcellular level.

## INTRODUCTION

Volume Electron Microscopy (vEM) techniques have emerged over the past 20 year addressing the intrinsic dimensional limitations of conventional EM, by enabling 3D reconstructions of comparatively large tissue volumes at nanometer-scale resolution. vEM encompasses methods based on transmission (TEM) and scanning electron microscopy (SEM) that generate image series which can be stacked to form a 3D digital representation of the original sample volume. Focused ion beam-scanning electron microscopy (FIB-SEM) is one of such vEM techniques, in which a focused gallium ion beam mills the block face at precisely controlled thicknesses, allowing substantially improved Z-resolution (Knott et al. 2008; Merchan-Perez et al., 2009). This approach removes material only from the sample surface rather than cutting the entire resin-embedded block, rendering the method only partially destructive and enabling tight process control (Knott et al. 2008; Merchán-Perez et al., 2009; Peddie et al. 2022). The availability of increasingly large vEM datasets, reaching petavoxel scales (Lichtman et al. 2014; Shapson-Coe et al. 2024), has driven the parallel development of advanced segmentation and 3D reconstruction tools, including both manual and automated pipelines, many of which leverage recent advances in machine learning (Xiao et al. 2018; Yin et al. 2020; Heinrich et al. 2021; Müller et al. 2024).

To date, vEM techniques have enabled the investigation of structure–function relationships across scales from organelles to the whole organism. In particular, they have proven to be of fundamental importance for the connectomics field, which faces the dual challenge of imaging neurons over sufficient volume to capture their full extent while maintaining the resolution required to visualize individual synapses as structural correlates of functional connections between cells (Peddie et al. 2022). vEM techniques have successfully used to uncover circuit-level architecture and layer-specific susceptibilities in both neurodevelopmental studies and neurological disorders (Beyer et al. 2013; Funke et al. 2019).

Here, we applied vEM to investigate the cytoarchitecture and ultrastructure of human brain organoids. Brain organoids are 3D tissue models derived from pluripotent stem cells and provide a unique opportunity to model brain development and neurological diseases, overcoming ethical limitations of human brain samples and animal models (Lancaster and Knoblich 2014; Maisumu et al. 2025).

To date, only a limited number of studies have applied vEM techniques to study the cytoarchitecture and ultrastructure of organoids (Jaros et al. 2017; Ronchi et al. 2021; D’Imprima et al. 2023; Jadav et al. 2023), and none has extended vEM approaches to micro-connectomic reconstruction or systematic mapping of synaptic architecture in brain organoids. Our work addressed this gap, by demonstrating the feasibility of such reconstructions and by providing a methodological framework for the future use of organoids as models of neurological or neurodevelopmental disorders or as platforms for therapeutic testing. Moreover, given the scarcity of literature on the topic, we compare two previously published embedding protocols (protocol 1: DeFelipe and Fairén 1993 and Cano-Astorga et al. (2024); protocol 2: Deerinck et al. 2010) to determine which is better suited for the processing of brain organoids, depending on the specific biological question.

To the best of our knowledge, this is the first study to establish a correlative vEM workflow enabling micro-connectomic reconstruction in human cortical organoids.

## MATERIALS AND METHODS

### Stem Cell culture and Organoid Differentiation

Healthy human induced Pluripotent Stem Cell (hiPSC) line WTSIi004 (male, age 35-39) was purchased from EBISC bank. hiPSCs were amplified and maintained using standard feeder-free culture protocols. In brief, hiPSC colonies were grown on Geltrex-coated plates (Gibco A1413201) in StemFlex medium (Thermo Fisher Scientific, A3349401) supplemented with Penicillin-Streptomycin (5,000 U/mL, Thermo Fisher Scientific), at 37°C in a humidified atmosphere containing 5% CO₂. Upon reaching 60-70% confluency, usually every 4-5 days, hiPSC colonies were passaged using Versene (Gibco 15040066) and treated with RevitaCell Supplement (Gibco A2644501) for 24 hours post-passage to support recovery. Growth medium was replaced every other day. hiPSC cultures were regularly checked for mycoplasma.

Dorsal Forebrain Organoids were generated using the protocol described in Sloan et al., 2018 (Sloan et al. 2018) with minor modifications. When hiPSC colonies reached around 1.5 mm in diameter, cells were detached by Accutase (Thermo Fisher Scientific, A1110501) and plated at day 0 (D0) into non-adherent U-shaped 96-well plates (Thermo Fisher Scientific, 174925) at the density of 20,000-25,000 cells in StemFlex medium supplemented with RevitaCell, 5 mM Dorsomorphin (DM) (Sigma-Aldrich, P5499) and 10mM SB-431542 (SB) (System/Tocris, 1614). At D2 and D4, the medium was exchanged and replaced with StemFlex supplemented with 5 mM DM and 10mM SB. From D6 to D25, organoids were grown in Neural Differentiation medium (NM), constituted by Neurobasal A medium (Thermo Fisher Scientific, 10888022), 2% B27 supplement without vitA (Thermo Fisher Scientific; 12587001), 1% GlutaMAX (Thermo Fisher Scientific 35050061), 1% Penicilin-Streptomycin (Thermo Fisher Scientific 15140122), and supplemented with 20ng/ml EGF and 20ng/ml FGF added freshly just before the use. Medium change was performed every other day. From D25 to D43, organoids were grown in NM supplemented with 20ng/ml BDNF (PeproTech, 450-02) and 20ng/ml NT3 (PeproTech, 450-03) added freshly just before the use and medium change was made every 2-3 days. From D43 onward, organoids were grown in NM without growth factors and medium change was performed every 4 days. After organoids grow larger (around D25-D30), they were moved to ultra-low-attachment 24-well plates (Thermo Fisher Scientific, 174930) and from D150 organoids were cultured with shaking so that the plates were moved to an orbital shaker.

Along this period (to 180 days), organoids growth was monitored. Images of each organoid during time were acquired by Incucyte or optical microscope and the area was measured by using an Arivis Cloud trained deep learning model as part of an image analysis pipeline in ZEN software from Zeiss (Carl Zeiss AG). Moreover, some organoids were fixed or collected at different time points to check for differentiation markers via immunofluorescence and western blot analysis.

### Immunofluorescence

At day 25, 44, 90, 180 cortical organoids were washed in PBS1x, fixed in Paraformaldehyde (PFA) 4% for two hours, washed again in PBS 1x and left in sucrose 30% to facilitate cryoprotection. After 48h, organoids were embedded in cryostat medium (Killik, Bio-Optica) and frozen at −80°C. Organoids were cut by cryostat (Leica Microsystems) into 20μm thin sections that were collected on 4% gelatin covered slides. Immunostaining was then performed. Slides were washed twice with PBS 1x to remove OCT/sucrose, permeabilized and blocked in PBS 1x, 10% NDS, 0.4% TritonX-100 for 1h at Room Temperature (RT) and then incubated with primary antibodies diluted in PBS 1x, 3% NDS, 0.4 Triton x-100 overnight at 4°C. Slides were washed three times in PBS 1x and then incubated with secondary antibodies and DAPI (to stain nuclei) diluted in PBS 1x, 3% NDS, 0.4 Triton x-100 for 1h at RT. Slides were washed again three times in PBS 1x, air-dried for 5-10 min, and coverslipped using mounting medium. Representative images of each primary antibody at different time points were acquired with Zeiss LSM 980 confocal microscope equipped with Airyscan.

### Western Blot

Fresh organoids were collected at day 90, 150 and 180 and hiPSCs were used as undifferentiated controls. For each time point, 6 organoids were pooled. Samples were lysed in ice-cold RIPA lysis buffer (Millipore, #20-188) supplemented with 1mM Na_3_VO_4_, 1mM PMSF, 1mM DTT and protease inhibitors (Roche Diagnostic, 11697498001). Samples were mechanically homogenized, kept on ice for 30min, then sonicated and centrifuged at 13,000rpm for 10 min at 4 °C. Supernatants were collected and concentration of proteins was determined using Bradford assay (Bio-Rad, 5000006). Samples were mixed with 4× LDS sample buffer (NP0007 and NP0009, Invitrogen), boiled at 95°C for 10 min, separated by SDS–polyacrylamide gel electrophoresis on 4–15% gradient gels (Bio-Rad, #4561086) and wet transferred overnight to PVDF (Bio-Rad, #1704274). Membranes were blocked with EveryBlot Blocking Buffer (Bio-Rad, #12010020) and incubated with the indicated primary and secondary antibodies diluted in 1%milk-PBS1x and then detected with Clarity Western ECL Substrate (Bio-Rad, #1705061) and Clarity Max Western ECL Substrate (Bio-Rad, #1705062).

### Organoid Fixation for EM analyses

Organoids were washed twice in PB 0.1M and fixed in PB 0.1M containing 2.5% glutaraldehyde and 2% formaldehyde for 2h at RT. Samples were then abundantly washed in PB 0.1M and stored at 4°C until further analysis.

### Araldite inclusion

For FIB-SEM imaging samples were prepared according to DeFelipe and Fairén (1993) and Cano-Astorga et al. (2024). Briefly, samples were treated with 1% OsO4 (Sigma, O5500, Germany), 0.1% potassium ferrocyanide (Probus, 23,345, Spain), and 0.003% CaCl_2_ in sodium cacodylate buffer (0.1 M) for 1 h at RT. They were then stained with 1% uranyl acetate (EMS, 8473, USA), dehydrated with a graded series of ethanol, cleared with acetone and flat-embedded in Araldite (TAAB, E021, UK).

Samples are then placed in the oven at 60° for 48 hours.

### Durcupan inclusion

Organoids were embedded in 4% agarose solution and, once the agarose had solidified, 100mm thick sections were cut on vibratome and stored in PB 0.1M at 4°C for later EM processing. This step was necessary to later allow heavy metal penetration and homogeneous contrast. We followed Deerinck et al. 2010. Compared to DeFelipe and Fairén (1993) and Cano-Astorga et al. (2024), this protocol involves additional steps in thiocarbohydrazide and Walton’s lead aspartate staining, leading to a stronger contrast of the membranes, and embedding in Durcupan resin. Samples were then placed in an oven at 60° for 48 hours.

### Semithin section collection and staining

Semithin sections (0.5 - 2 µm thick) were obtained using an ultramicrotome (Leica MZ6) equipped with a 6 mm Diatome Histo diamond knife. Sections are then collected on water drops on glass slides previously coated with silicone and an araldite layer, which facilitated their subsequent detachment. In some cases, the slides were additionally sputter-coated with heavy metals to enable SEM observation, following the preparation protocol described by Rodríguez et al. (2018).

Sections were then stained with 1% toluidine blue (Merck, 115,930, Germany) in 1% sodium borate (Panreac, 141,644, Spain), examined under light microscope and photographed. The sections of interest for SEM imaging (**Figure 1A**) or TEM were then trimmed with a single-edged cutter blade and the araldite piece containing the selected section was detached using the underlying silicone layer.

**Figure 1:**
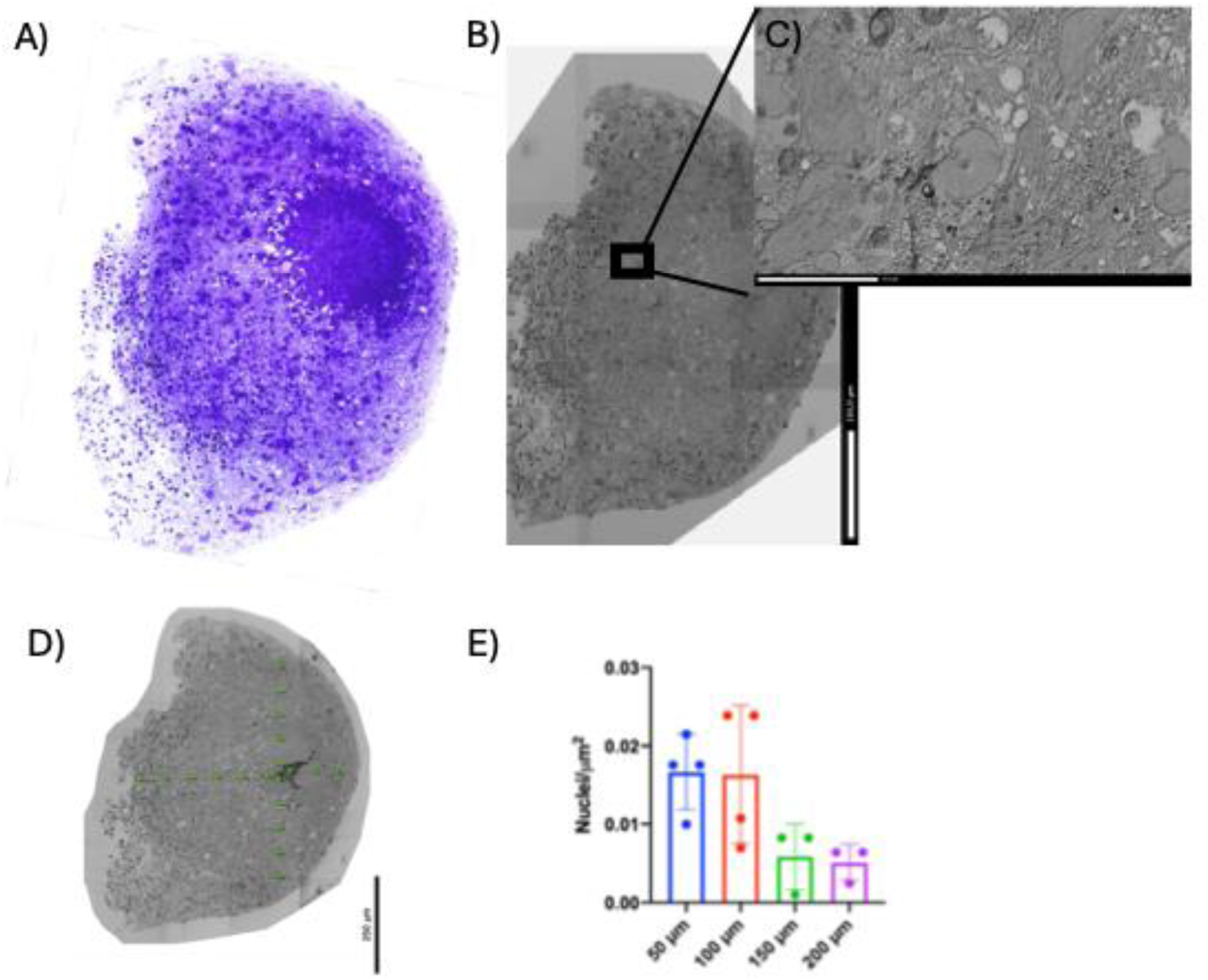
Organoid region characterization. (A) Toluidine blue staining of the last semithin section before block-face (B) Mosaic acquisition obtained with Atlas 3D software and visualized at the original resolution in the AtlasViewer program with SE2 detector. (C) Inset of the mosaic acquisition (resolution 20 nm). (D) Flag markers (green dots) positioned from the rosette outward, indicating the regions of interest (ROIs) where the nuclei were counted. (E) Quantification of nuclear density within the identified ROIs. The scalebars are shown in each image.

### SEM imaging of semithin sections

The section selected (0.5 µm thick) for subsequent imaging was the last one, adjacent to the before block-face.

Each section was then mounted onto a SEM specimen stub, using conductive carbon adhesive tab (Electron Microscopy Sciences, 77825-09) and silver paint (Electron Microscopy Sciences, 12630). After drying for 24 h in a vacuum desiccator, samples were imaged using a field emission SEM (Gemini II column, Zeiss, Germany).

Mosaic acquisition of the entire organoid surface was obtained using Atlas 3D software and at naive resolution with AtlasViewer (**Figure 1B**).

### FIB-SEM imaging

To perform FIB-SEM imaging, the blocks were glued onto a sample stub using conductive carbon adhesive tab. All surfaces, except the top surface, were covered with silver paint to prevent resin charging. The stubs were then sputter-coated (with Emitech K575X, Quorum Emitech, Ashford, Kent, UK) with gold/palladium for 60 seconds to facilitate charge dissipation. We carried out FIB-SEM imaging (Crossbeam® 40 electron microscope, Carl Zeiss NTS GmbH, Oberkochen, Germany) on the “mirror” side opposite to the last semithin section from which the mosaic was taken. In this way the mosaic can be used as a reference to distinguish the different regions of the organoid (including rosettes, and neuropil-like regions) (**Figure 2B**).

**Figure 2:**
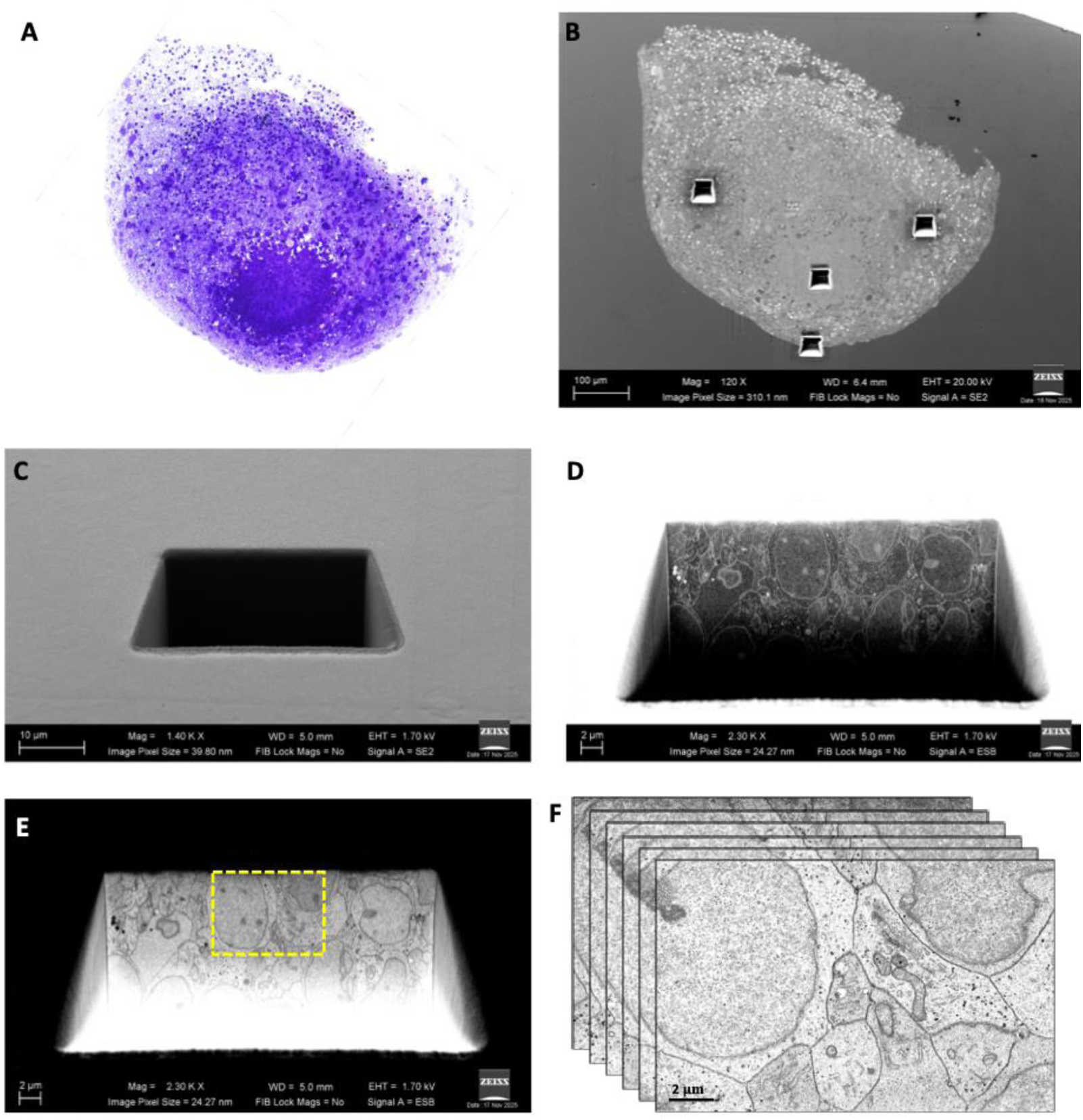
FIB-SEM imaging workflow (A) Toluidine staining of the last semithin section before block-face (B) The block face with the organoid surface is visible. On the surface three peripheral trenches and a central one can be observed. Image stacks analyzed in this study were acquired from these regions (C) Magnification of a trench observed with secondary electrons (SE2) (D) The surface exposed by FIB milling is imaged by backscattered electrons (ESB) (E) The same image as in C is inverted. The yellow inset indicates the regions from which the image series was obtained (F) Representative illustration of the image series obtained from the central trench. The scalebars are shown in each image.

This technique combines a high-resolution field-emission SEM column with a focused gallium ion beam (FIB), which permits removal of thin layers of material from the sample surface at the nanometer scale. A first trench (**Figure 2C-E**) was obtained with a current of 7 nA, then the inner surface visible in the trench was cleaned with a current of 750 pA. The FIB removed a thin layer of material (20 nm thick) and the newly exposed surface was imaged by the SEM (1.7 kV) using the backscattered electron detector (ESB) (Knott et al. 2008; Merchan-Pérez 2009; Blazquez-Llorca et al. 2013). In this way, long series of images were acquired from selected regions of the 3D samples (Merchan-Pérez 2009) (**Figure 2F**). Resolution in the *z*-axis (section thickness) was 20 nm, the image size was 2048 × 1536 pixels, and the in-plane resolution 5 nm/pixel. Each image stack comprised approximately 350 serial sections.

### Stacks alignment and visualization

The image stack alignment (registration) was performed using Fiji (http://fiji.sc). The “Register Virtual Stack Slices” plug-in was applied using a rigid registration method (translation only, no rotation) to avoid deformation of single sections.

We then applied a Gaussian Blur filter with Fiji to eliminate noisy pixels and facilitate the following segmentation process.

### EspINA software

EspINA software was used for the automated segmentation and counting of synapses in the reconstructed 3D volume (Morales et al. 2011: https://cajalbbp.csic.es/espina-2/) and for the tracing of neurite “skeletons” as described in Turegano-Lopez et al. (2024).

The synaptic density was determined by quantifying the segmented synapses within a predefined 3D counting frame of known volume (Merchan-Pérez 2009). An unbiased stereological approach was employed, using a 3D counting box covering most of the examined tissue and delimited by three inclusion and three exclusion planes (Merchan-Pérez 2009). The EspINA software identifies the segmentations within this volume, counting those in contact with inclusion planes while excluding those intersecting the exclusion planes (Morales et al. 2011). The segmentation process applies a gray-level threshold to isolate voxels corresponding to pre- and postsynaptic densities, visible as electron-dense regions under the electron microscope (Morales et al. 2013).

Moreover, to reconstruct the connectivity of the organoid, we took advantage of a tracing tool available in EspINA (Turegano-Lopez et al. 2024). This allows to follow the trajectory of each neurite throughout the image series and to reconstruct their branching (**Figure 3A**). Since synapse segmentation is performed first, the post-synaptic element was classified as a dendrite, while the pre-synaptic one, in which synaptic vesicles were also visible, was classified as an axon. Fibers that did not make synapse within the imaged volume were classified as neurites.

**Figure 3:**
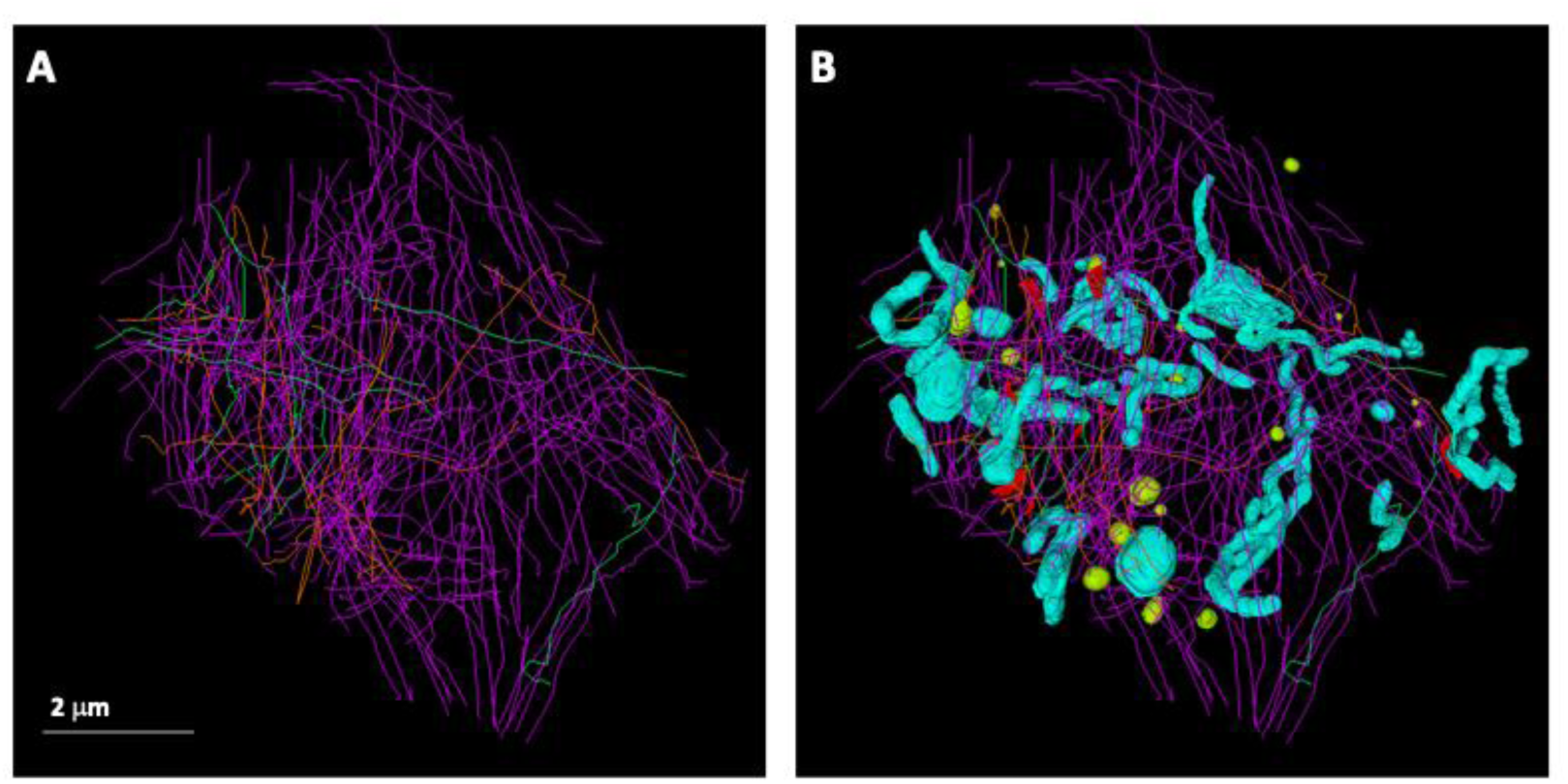
3D reconstruction using EspINA software of neurites, synaptic elements and organelles from a stack of 370 images of a 3-month-old organoid. (A) Skeleton reconstruction showing traced neurites (magenta), axons (green) and dendrites (orange). (B) Same reconstruction as in (A) with synapses (red), multivesicular bodies (yellow) and mitochondria (cyan). All reconstructions were generated from a FIB-SEM datasets obtained as previously described.

EspINA software also gives quantitative data (examples are given in **Figure 4**) for each skeleton, such as length and the number of synapse made/received. Moreover, EspINA can automatically extract synapse apposition surface (SAS) from the segmentation. This measure has a functional relevance, since this area correlates with the probability of neurotransmitter release and with receptor abundance.

**Figure 4:**
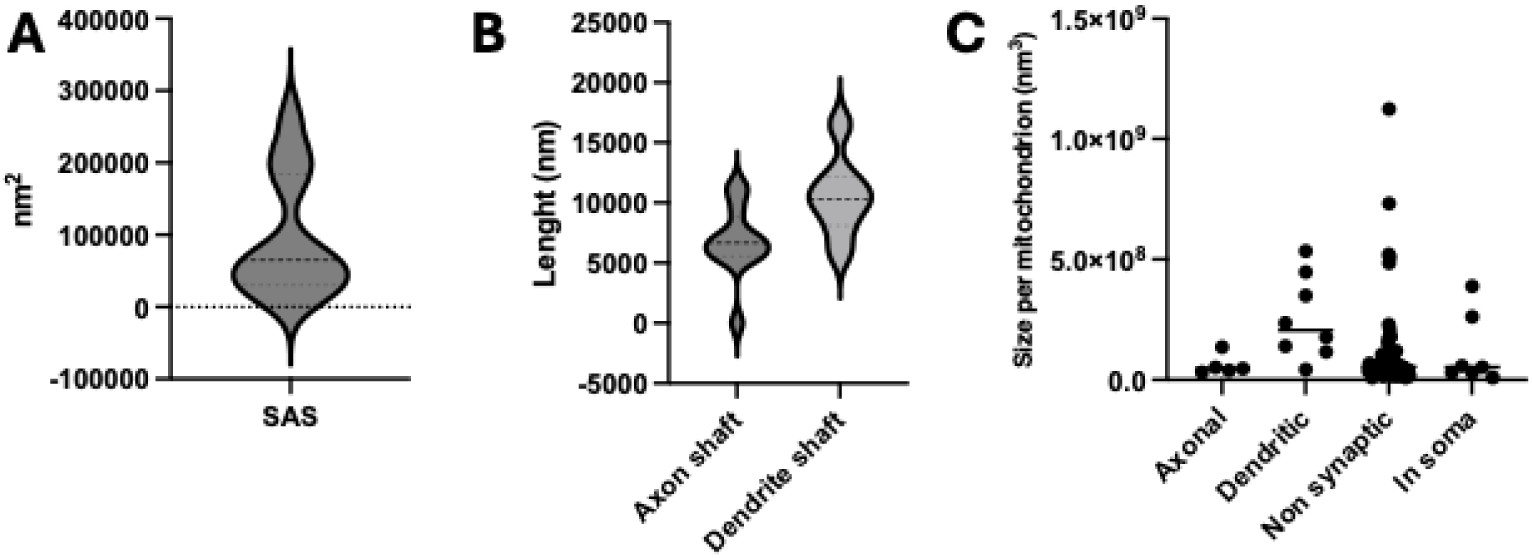
Quantitative analyses obtained by EspINA software after neurite, synapse and mitochondria segmentation. One image stack from a 3-month old human cortical organoid was analyzed using EspINA software as a proof of concept, illustrating the type of quantitative data that can be obtained. (A) Violin plot representing medium synaptic apposition surface (SAS) and its distribution. (B) Violin plot showing the average axon and dendrite length within the analyzed stack. (C) Mitochondrial size distribution in each compartment: axonal, dendritic, non-synaptic fibers, and soma.

Finally, organelles can be manually segmented in 3D using the “Segment” tool in EspINA (**Figure 3B**), allowing the measurement of their dimensions. Additional quantitative data can also be obtained, such as the Volume % occupied by a class of organelles or their distribution across different compartments (axon, dendrite, neurite), being the skeleton available.

### TEM

Ultrathin sections can be obtained either from the same block used for FIB-SEM imaging (after trimming it to the appropriate size and a trapezoid shape) or from semithin sections (1-2 µm thick). Semithin sections were stained with toluidine blue and examined under a light microscope. The section of interest was then mounted on top of a resin cone, and the region of interest trimmed again into a trapezoid (**Figure 5A**). This newly trimmed block face can be re-imaged under the light microscope. Ultrathin sections (50–70 nm thick) were cut with a diamond knife at the ultramicrotome (Leica MZ6). The ultrathin sections were collected on formvar-coated grids. Samples prepared according to DeFelipe and Fairèn (1993) and Cano-Astorga et al. (2024) required further grid staining with uranyl acetate and lead citrate, while this step was not necessary for samples included with the method of Deerinck et al. (2010). Digital pictures were captured at different magnifications using a JEOL JEM-1011 transmission electron microscope (JEOLLtd., Tokyo, Japan) equipped with a digitalizing image system (SC1000 ORIUS, 11megapixel; Gatan, Pleasanton, CA).

**Figure 5:**
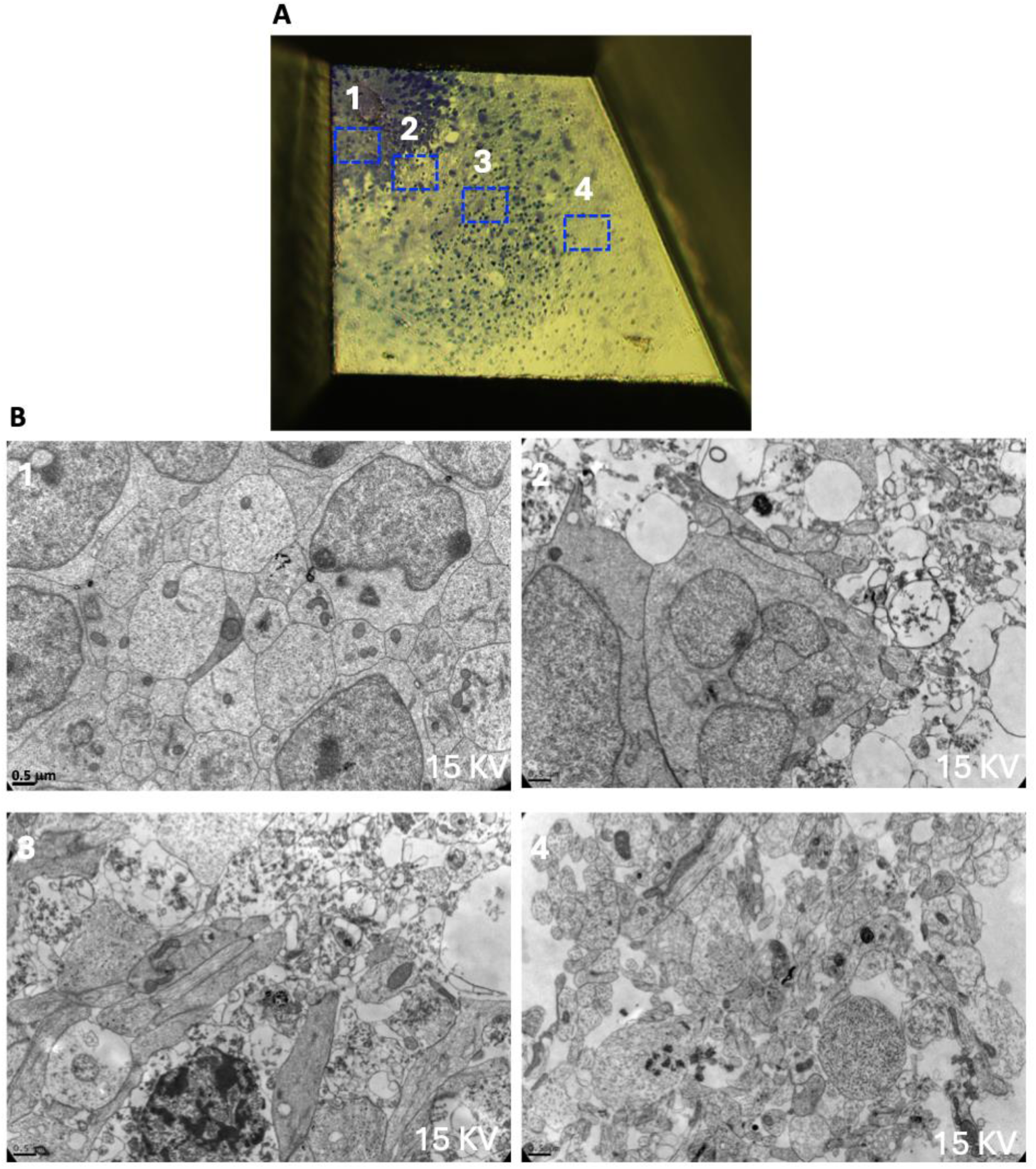
TEM imaging. (A) Block face from which ultrathin section are cut. A semithin section stained with toluidine is trimmed to a smaller trapezoid shape. Red insets indicate the different regions (B) Corresponding TEM images acquired from the regions indicated in (A).

### Statistical analyses

Stereological counts of nuclei density (**Figure 1E**), data obtained from EspINA software (**Figure 4**), organoid area measurements and western blot analyses (**Figure S1**) were plotted and analyzed using GraphPad Prism 9.0. Organoid area at the different time points were compared using One-way Anova, with Turkey’s multiple comparisons test. For EM data statistical analysis was not performed, as long as the data shown in Figure 1E and 4 are only representative of the data that can be obtained from stereological count, neurite tracing and segmentation.

## RESULTS

### Correlative light and electron microscopy of semithin sections: imaging and characterization of organoid regions

Cortical organoids (specifically dorsal forebrain organoids) were generated following the guided protocol developed by Sloan et al., (2018) with minor modifications as described in the Materials and Methods section. Organoids were maintained in culture up to 180 days and showed a steady, significant growth and maturation. At the initial stages (up to 60 days) we observed a remarkable expansion in their dimensions, which stabilized at later stages (**Figure S1A**). Presumably intermediate and later stages reflected ongoing maturation and neuronal differentiation of organoids. Indeed, by immunofluorescence and western blot analyses, performed at different time points of growth, we detected the expression of markers typical for neural stem cells (SOX2, PAX6), early after organoid formation; neurons (MAP2), already at 25 days; cortical neurons of layer II/III (SatB2) and V (CTIP2) and astrocytes (S100b, GFAP), increasingly at later stages (180 days). These data confirm the development of a high degree of tissue organization in our cortical organoids (**Fig S1B-C**).

For EM analysis, we choose organoids at 3 months of differentiation, as they contained both immature and mature neurons, allowing a more comprehensive analysis of organoid architecture. For EM imaging, we tested and compared two different protocols: DeFelipe and Fairén, 1993/Cano-Astorga et al. (2024) and Deerinck et al (2010) protocol. Semithin section of 500 nm were collected on sputter-coated microscope glass slide covered with silicone, Araldite (as described in Rodríguez et al. 2018). The sections were then dried and stained with toluidine blue (**Figure 1A** and **2A**).

These sections were examined and viewed under a light microscope. Toluidine blue staining revealed the structural heterogeneity of the organoids. Indeed, as described by Paşca et al. 2015, cortical organoids display ventricular zone-like structures, also referred as rosettes, containing proliferating cells. The rosette can be appreciated in semithin sections as nuclei-dense regions, with a radially organized, concentric arrangement of neural progenitor cells.

The section of interest was then trimmed from the glass slide and mounted onto a SEM specimen stub. The Atlas3D software was used to obtain mosaic acquisitions of the whole section using the SE2 detector with a resolution of 20 nm (**Figure 1B-C**). Smaller regions of interest can be imaged at higher resolution and/or using different detectors (such as ESB). Ideally, the last section cut immediately before the block face should be used for this step, as it provides a 2D mirror image of the first block-face section: this enables accurate spatial mapping before eventually moving to more specific volume EM imaging of the target regions by FIB-SEM. This workflow was best suited on organoids processed using the DeFelipe and Fairén (1993)/Cano-Astorga et al. (2024) inclusion method, as the lower contrast and the use of Araldite resin provide optimal toluidine blue staining, compared to a stronger heavy metal staining and Durcupan resin (a comparison of the advantages and limitations of each technique is shown in **Table 1**).

**Table 1:** the table summarizes advantages and limitations identified for each inclusion protocol.

|  | Deerinck’s protocol (2010) + Durcupan | DeFelipe and Fairén (1993) / Cano-Astorga et al. (2024) protocol + Araldite |
| --- | --- | --- |
| PROS | • Does not necessitate TEM grid staining<br>• Higher membrane contrast may facilitate automatic segmentation | • High quality toluidine blue staining of semithin section, allowing easier region identification<br>• Reliable post-synaptic density identification |
| CONS | • Poorer toluidine blue staining of semithin sections<br>• Difficult identification of post-synaptic density | • Requires TEM grid staining<br>• Lower membrane contrast |

Starting from the rosette, we performed a stereological count of nuclei density (**Figure 1D-E**). In detail, using the mosaic image, regions were defined at 50 μm intervals moving radially outward from the center of the rosette. Then, in EspINA software, in the labeled regions we counted the nuclei within a counting frame of 3,500×4,000 nm, with 2 inclusion and 2 exclusion borders. The nuclear density was calculated as nuclei/μm^2^ and averaged across the four radial directions at each distance from the reference rosette. As expected, the number of nuclei decreased with increasing distance from the rosette.

### Synapse/organelle segmentation and neurite tracing by EspINA software

We also aimed to assess in cortical organoids the feasibility of applying the same analyses previously performed in the mouse somatosensory cortex by Turegano-Lopez et al., 2024 and, ultimately, to study the connectome of the organoid at the ultrastructural level. The mosaic imaging and the stereological count of the nuclei (**Figure 1D-E**) provided a spatial map to guide trench opening and FIB-SEM imaging (**Figure 2**) in regions most likely to contain neuropil-like areas, characterized by low nuclear density and a high density of neurites. Therefore, for each time point, we placed 3 trenches in peripheral regions, near the organoid borders (**Figure 2B**). For this analysis, the FIB-SEM is particularly suitable as it allows precise sampling in distinct organoid regions, partially overcoming the problem of the high heterogeneity of organoid tissue (Yoon et al. 2019).

We then segmented synapses and traced neurites by EspINA software, as described in Turegano-Lopez et al. (2024) (**Figure 3**). This analysis enabled the extraction of quantitative parameters, including synaptic apposition surface (SAS) (95,862 nm^2^ in our stack) and neurite length (axon shaft length: 6,827 nm; dendrite shaft length: 10,530 nm) (**Figure 4A-B**).

This analysis yielded the most accurate results when FIB-SEM was applied to samples embedded as in DeFelipe and Fairén (1993)/Cano-Astorga et al. (2024). This is because FIB-SEM allows thinner cutting and thus higher resolution in z-axis, compared to SBF-SEM, which is critical for accurate synapse reconstruction. Moreover, the excessive contrast, due to aspartate solution in Deerinck’s protocol (2010), prevents a precise visualization of the postsynaptic density. The analysis was feasible also in samples imaged in FIB-SEM following inclusion with Deerinck’s protocol.

Moreover, as a feasibility test, in one image stack from an araldite-embedded sample, we performed 3D segmentation also of all the mitochondria and multivesicular bodies (MVBs). Thanks to the previous segmentation of synapses and neurite tracing, it was possible to obtain quantitative data, such as volume fraction occupied by mitochondria or MVB and their abundance in each compartment (axon, dendrite, non-synaptic fibers) (**Figure 4C**).

### TEM validation of FIB-SEM ultrastructural findings

With FIB-SEM, it is possible to achieve a resolution comparable to that obtained with TEM. However, in vEM imaging, a balance between resolution and acquisition time must be considered. For this reason, to obtain further ultrastructural details, we cut ultrathin sections and performed TEM imaging. As described, ultrathin sections can be obtained either from the same block used for FIB-SEM imaging or from semithin sections (**Figure 5A**). Using the toluidine blue image as a map (**Figure 4A**), we imaged different regions from the rosette outward, at high resolution (**Figure 5B**). This approach confirmed the presence of synapses (previously identified with FIB-SEM), and the possibility to recognize synaptic vesicles, a well-defined postsynaptic density, and closely apposed pre- and postsynaptic membranes (**Figure 6**). Moreover, the high degree of cellular packing within the rosette was evident. Indeed, in this region, the intercellular space between adjacent membranes was minimal, and in some areas formed structures suggestive of adherens and tight junctions (**Figure 7E-F**). Intracellular organelles and degenerating structures can also be observed (**Figure 7A-B**).

**Figure 6:**
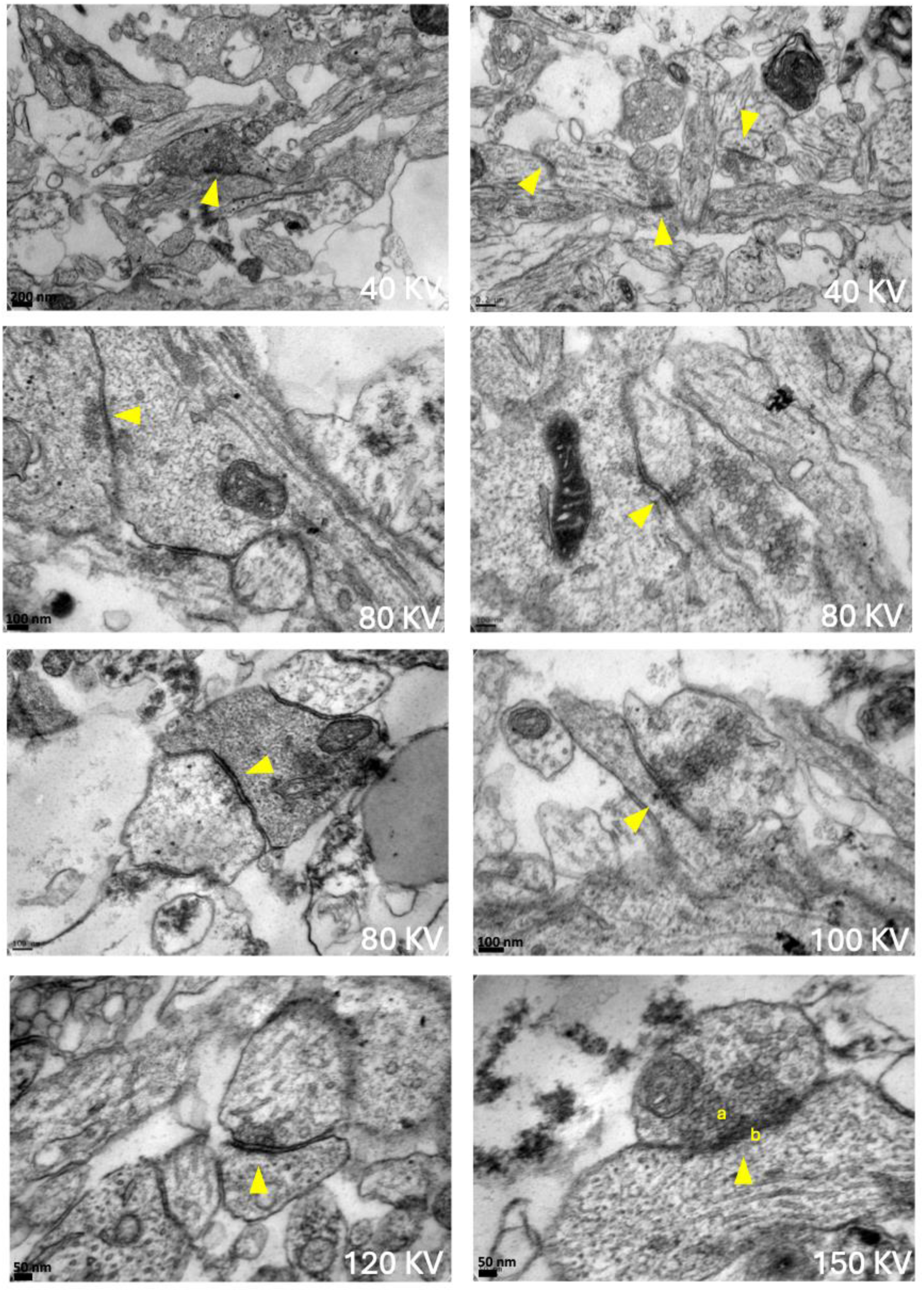
Synapses are indicated with yellow arrows. In the last image, the one at higher magnification, the pre-synaptic zone, rich in synaptic vesicles is indicated with (a), the post synaptic density with (b).

**Figure 7:**
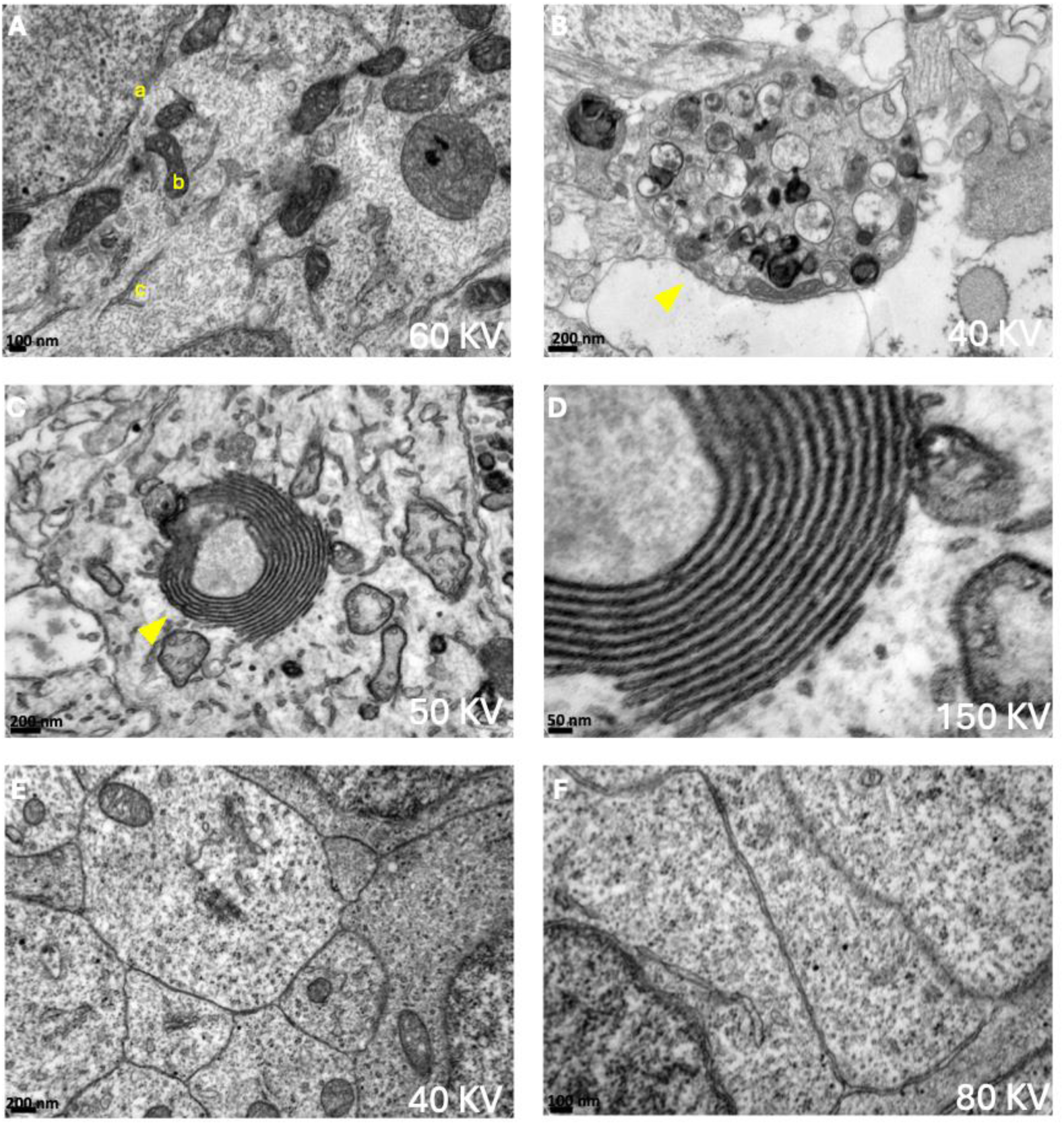
High resolution imaging of different ultrastructural structures. (A) Different organelles are visible, such as part of a nucleus (a) with double nuclear membrane, mitochondria (b) and endoplasmic reticulum (c). (B) Dystrophic neurite containing autophagic vacuoles and degenerating structures. (C) and (D) Examples of organized tubular structures of endoplasmic reticulum. (E) and (F) Tightly packed cells within the rosette; adjacent cell membranes display minimal intercellular space.

**Figure 8:**
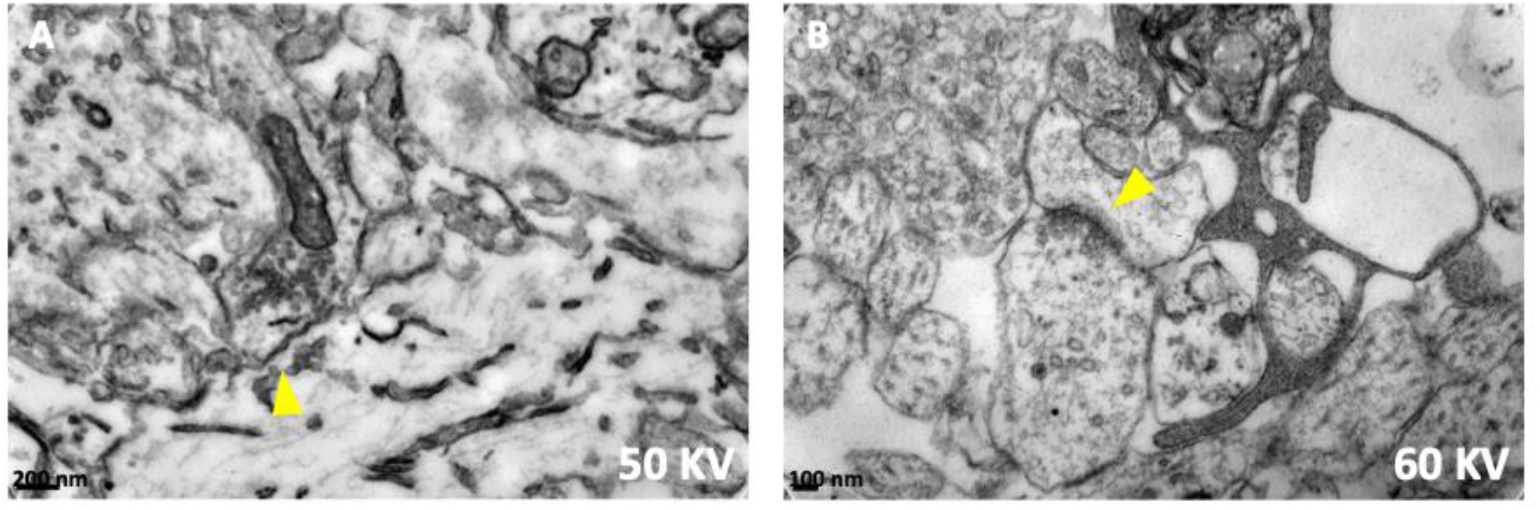
Comparison between Deerinck’s protocol (2010) (A), and DeFelipe and Fairén (1993)/Cano-Astorga et al. (2024) protocol (B). Synapses are indicated with yellow arrows. In this Figure the different level of contrast can be appreciated.

## DISCUSSION

In this study, we established a comprehensive workflow for applying vEM to the analysis of human cortical organoids, integrating correlative light and electron microscopy, FIB-SEM imaging, and TEM validation. We compared two widely used but conceptually divergent approaches to electron microscopy preparation, because they represent opposite ends of the EM contrast spectrum. We showed that the DeFelipe and Fairén (1993)/Cano-Astorga et al. (2024) method provides superior compatibility with toluidine-based semithin sectioning and yields optimal contrast for synapse segmentation and neurite tracing. This is of crucial, as synapse classification relies on the relative electron density and thickness of the postsynaptic density, which distinguishes asymmetric (typically excitatory) from symmetric (typically inhibitory) synapses (Peters et al. 1991; Peters and Palay 1996). However, the validity of this classification —typically applied to native tissue—should be further confirmed in organoids by immunostaining analysis (e.g. for GABAergic and glutamatergic markers). Conversely, Deerinck’s protocol (2010) offers enhanced membrane contrast, but introduces limitations for postsynaptic density visualization. Related to this, a higher contrast could help in implementing automatic segmentation pipelines, which would also enable larger volume reconstruction as has been recently done by Shapson-Coe and colleagues with a petavoxel fragment of human cerebral cortex (Shapson-Coe et al. 2024). The main pros and cons we identified for each methods were summarize in **Table 1**.

We further demonstrated that FIB-SEM imaging of peripheral, neuropil-like regions enables accurate 3D reconstruction of synapses, neurites, and intracellular organelles, allowing quantitative assessment of synaptic apposition surfaces, neurite trajectories, and organelle distribution within defined cellular compartments.

Collectively, our results confirm that vEM is fully applicable to cortical organoids (and more broadly to brain organoids) and can map their micro-connectome with nanometer resolution. This methodological framework provides a scalable foundation for future studies aimed at characterizing organoid ultrastructure during development and at different maturation stages, comparing disease-specific organoid models, and evaluating therapeutic interventions at the level of synaptic and subcellular levels. Indeed, degenerating structures such as dystrophic neurites are readily identifiable (**Figure 7B**) and expected even in healthy organoids (**Figure 6**). Their presence and spatial distribution could be compared across different experimental conditions, relating them to specific disease-related mutations or other altered pathogenic mechanisms. Similarly, this approach enables detailed 3D morphometric analysis of subcellular organelles (**Figure 3B, 4C and 7A)**, such as mitochondria, —whose dysfunction is central to many neurodegenerative diseases. For example, in Alzheimer’s disease, altered mitochondrial morphology (fragmented cristae, reduced size, dense material accumulation) reflects stress impaired bioenergetics and increased oxidative stress that may contribute to disease progression (Vaillant-Beuchot et al. 2021). Similar dysregulation of mitochondrial dynamics —excessive fission, impaired fusion, disrupted axonal/dendritic transport— has been implicated in Parkinson’s disease, motor neuron diseases and other neurodegenerative conditions, linking morphological changes to synaptic dysfunction and neuronal loss (Stanga et al. 2020). These disease-associated alterations underscore the value of FIB-SEM and TEM for quantitative mitochondrial comparisons across compartments and conditions in organoid models. Analyzing different regions within the same organoid (i.e. 3 peripheral and one central) and enlarging the sample size are critical factors for enabling meaningful comparisons across different differentiation time points and for the EM-based characterization pathological organoid models. This approach is particularly important given the high intrinsic variability that characterizes these in vitro systems (Yoon et al. 2019).

In this study, we used EspINA due to its extensive use in the literature for FIB-SEM stack analyses of cortical tissue (e.g. Morales et al. 2011; Domínguez-Álvaro et al. 2018, 2019; Cano-Astorga et al. 2023) and its free availability. However, other open-source software tools could also be applied for connectomics and segmentation in organoids, including platforms for manual and semi-automated reconstructions such as TrakEM2, VAST, Microscopy Image Browser and CATMAID (Kievits et al. 2022).

Finally, current and future advances in machine learning are expected to greatly accelerate the analysis of 3D vEM datasets, a phase that remains highly time-consuming with conventional manual or semi-automated approaches.

Overall, to our knowledge, this study presents the first correlative vEM workflow enabling micro-connectome reconstruction in human cortical organoids. This approach offers a powerful tool for future investigations of neurodevelopmental processes and neurological disease modeling.

## ACKNOWLEDGEMENTS

SD’s research has been conducted during and with the support of the Italian national inter-university PhD course in Sustainable Development and Climate Change (link: www.phd-sdc.it).

## AUTHOR CONTRIBUTION

Conceptualization: MTL, AV, JD, MB; Methodology: SD, AM, MTL, LBL, RS, AV, JD, MB; Investigation and data curation: SD, AM, MTL, LBL, AMP, RS; Visualization: SD, AM, AMP, MTL; Writing – Original Draft: SD, AM, MB; Writing – Review & Editing: All authors; Supervision: MTL, AV, JD, MB; Funding Acquisition: LBL, AV, JD, MB.

## SUPPLEMENTARY FIGURE

**Supplementary Figure 1.**
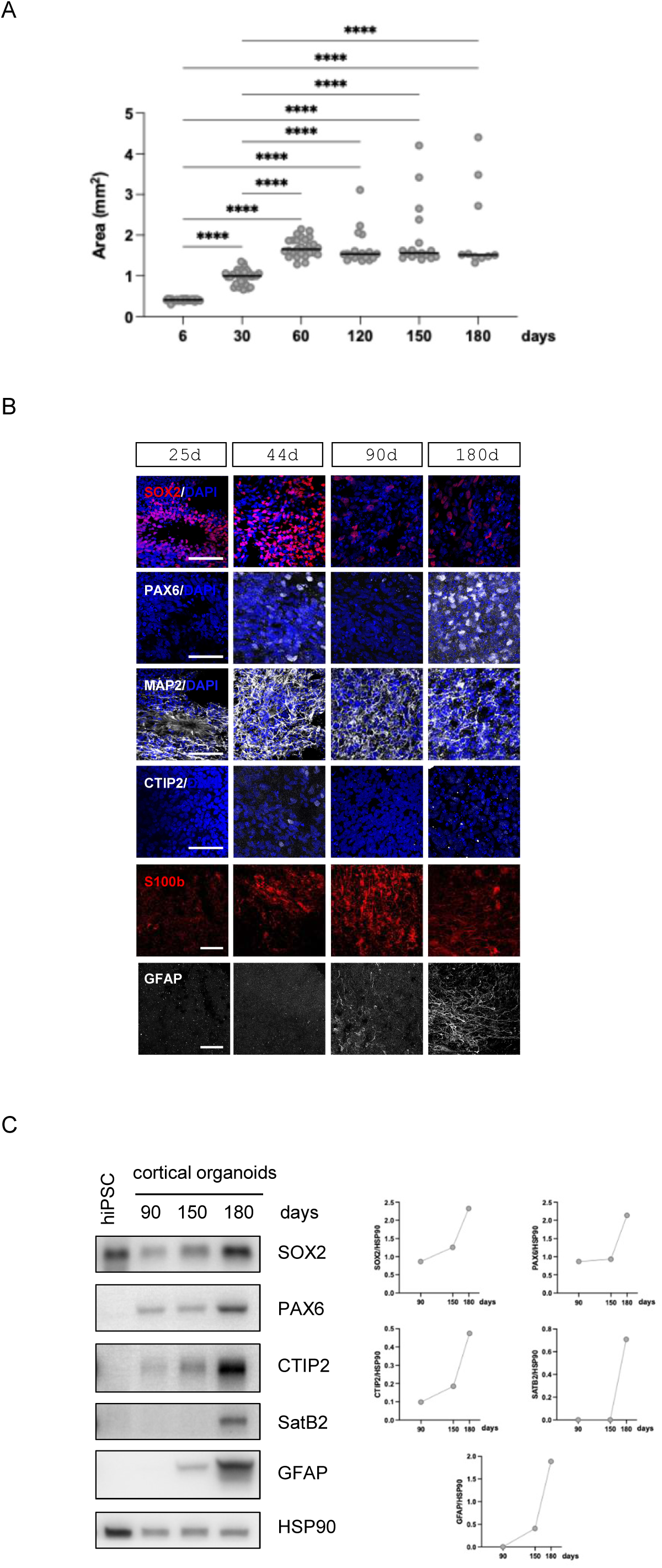
Human iPSCs-derived cortical organoids show mature morphology and neuronal and glial markers expression. A) The area of organoids at the indicated time points was reported in the graph and shows organoids growth. Median is shown as solid black line. 6days and 30days n=32; 120days n=18; 150days n=17; 180days n=10. ****p<0.0001 (One-way ANOVA, Turkey’s multiple comparisons test). (B,C) Representative images of immunofluorescence (B) and western blot (C) analyses performed to monitor maturation of cortical organoids. Neural progenitor cells (SOX2, PAX6), neuron (MAP2), cortical neurons of layer II/III (SatB2) and V (CTIP2); astrocytes (S100b, GFAP). (B) DAPI was used to stain nuclei. Scale bar: 50mm. (C) For the WB, HSP90 was used as loading control. For each time point 6 organoids were pooled together.

